# Parechovirus-3 infection disrupts immunometabolism and leads to glutamate excitotoxicity in neural organoids

**DOI:** 10.1101/2024.09.10.611955

**Authors:** Pamela E. Capendale, Anoop T. Ambikan, Inés García-Rodríguez, Renata Vieira de Sá, Dasja Pajkrt, Katja C. Wolthers, Ujjwal Neogi, Adithya Sridhar

**Affiliations:** OrganoVIR Labs, Emma Children’s Hospital, Department of Pediatric Infectious Diseases, Amsterdam UMC, location Academic Medical Center, Amsterdam Institute for Infection and Immunity, Amsterdam Institute for Reproduction and Development, University of Amsterdam, Meibergdreef 9, 1105 AZ Amsterdam, The Netherlands; OrganoVIR Labs, Department of Medical Microbiology, Amsterdam UMC, location Academic Medical Center, Amsterdam Institute for Infection and Immunity, University of Amsterdam, Meibergdreef 9, 1105 AZ Amsterdam, The Netherlands; The Systems Virology Lab, Division of Clinical Microbiology, Department of Laboratory Medicine, Karolinska Institutet, ANA Futura, Campus Flemingsberg, Stockholm, Sweden; UniQure Biopharma B.V., Department of Research & Development, Paasheuvelweg 25A, 1105 BE Amsterdam, The Netherlands; Emma Center for Personalized Medicine, Amsterdam UMC, Amsterdam, the Netherlands

**Keywords:** Neural Organoids, Immunometabolism, Parechovirus, Quantitative Proteomics, Central Nervous System

## Abstract

*Parechovirus ahumpari 3 (HPeV-3)*, is among the main agents causing severe neonatal neurological infections such as encephalitis and meningitis. However, the underlying molecular mechanisms and changes to the host cellular landscape leading to neurological disease has been understudied. Through quantitative proteomic analysis of HPeV-3 infected neural organoids, we identified unique metabolic changes following HPeV-3 infection that indicate immunometabolic dysregulation. Protein and pathway analyses showed significant alterations in neurotransmission and potentially, neuronal excitotoxicity. Elevated levels of extracellular glutamate, lactate dehydrogenase (LDH), and neurofilament light (NfL) confirmed glutamate excitotoxicity to be a key mechanism contributing to neuronal toxicity in HPeV-3 infection and can lead to apoptosis induced by caspase signaling. These insights are pivotal in delineating the metabolic landscape following severe HPeV-3 CNS infection and may identify potential host targets for therapeutic interventions.

## Introduction

In the absence of a self-contained metabolism, viruses hijack host proteins and pathways to facilitate viral replication(1). In response, host cells deploy antiviral mechanisms, such as interferon signaling and release of other cytokines, resulting in an inflammatory microenvironment. Due to the additional energetic demands imposed by the viral replication and the consequent antiviral responses, metabolic alterations are observed in host cells, a phenomenon referred to as metabolic reprogramming(2-4). In recent years, there has been increasing recognition of metabolic reprogramming due to viral infections, in addition to immune responses, as an important factor in disease progression, severity, and duration(5).

Recently, we reported heightened innate inflammatory responses as a potential cause for differences in neurologic disease caused by different *Parechovirus ahumpari (HPeV)*, previously *Parechovirus A*, genotypes that belong to the *Picornaviridae* family(6). However, the alterations in the cellular metabolic landscape following HPeV infections have not yet been studied. Other picornaviruses have been shown to exert a significant effect on the host cell metabolism in a range of different tissues *in vitro* and *in vivo*. This includes shutting off the immune response and regulating glucose, glutamine, lipid, and nucleotide metabolism to maximize the number of viral progenies(7-11). Additionally, they can extend the lifespan of infected cells to ensure the completion of the viral replication cycle(12, 13). Insights into the early metabolic footprints can help to identify factors contributing to long term disease outcome(5, 14). Understanding metabolic reprogramming following HPeV-3 infection is important as a large proportion of young children (27% of children younger than three years) infected with HPeV-3 show neurological sequelae and neurodevelopmental delay in long-term follow-up(15-18).

The specific metabolic changes that occur after viral infection depend on the virus and tissue type, owing to the unique metabolic environment of different tissues(1, 19, 20). The central nervous system (CNS), in particular, has a distinct metabolic footprint in which, amongst others, astrocytes and neurons are highly interactive. A disruption of metabolic homeostasis between neurons and astrocytes can initiate glutamate excitotoxicity and lead to apoptosis induced by increased reactive oxygen species (ROS) levels(21). Glutamate excitotoxicity is a crucial pathological factor in acute and chronic neurodegenerative conditions(22-24). Viral infections can also disrupt this equilibrium, resulting in neuronal damage(25). A previous study using primary mouse astrocytes and neurons showed that herpes simplex virus type 1 (HSV-1) infection resulted in an increased amount of extracellular transmitters such as glutamate(26). Other RNA viruses, *e*.*g*. human immunodeficiency virus (HIV-1), are known to cause dysregulation of the excitatory neurotransmitter release, contributing to neuronal and glial dysfunction. The correlation between this glutamate dysfunction and HIV-1-associated neurocognitive disorder (HAND) has been widely studied (27-29). However, no studies have been conducted on HPeV, leaving a gap in understanding the impact of HPeV infection on CNS metabolic homeostasis.

To this end, we investigated the impact of HPeV infection on the CNS metabolic footprint using stem cell-derived neural organoids. By using our quantitative proteomics dataset, we identified differences in metabolic changes induced by HPeV-1 and HPeV-3 infection that link to neuropathological features unique for HPeV-3 infection. Involvement of glutamate excitotoxicity was confirmed by measuring extracellular of glutamate and LDH levels. We validated our findings using functional assays in human induced pluripotent stem cell-derived (hiPSC) neural-astrocyte co-cultures.

## Results

### Proteome analysis of HPeV-infected neural organoids shows proteins, pathways, and metabolic dysregulation unique for HPeV-3

We previously published a proteome dataset highlighting differences in immune response in neural organoids infected with HPeV-1 and HPeV-3(6) and the relation to their clinical presentation (figure 1A). By analyzing this dataset with a focus towards the metabolic processes, we aimed to obtain insights into metabolic reprogramming in neural organoids following HPeV infection. Consistent with the previous analysis(6), differentially abundant proteins (DAPs) were immune-related in neural organoids between HPeV-1 and HPeV-3, compared to mock-infected organoids (figure 1A-C). To distinguish factors specifically contributing to HPeV neuropathogenicity, we directly compared DAPs between HPeV-1 and HPeV-3 infection. We identified 19 proteins (HTRA1, CLU, PLCG1, PRSS22, IGSF3, PPP1R1B, PSAT1, PDLIM3, PTDSS1, MTM1, USP5, CHMP3, MOB4, PLPPR3, GOSR2, TSR1, HOMOX2, DTD1, SST) to be uniquely abundant in HPeV-3 infected organoids (figure 1D).

**Figure 1.**
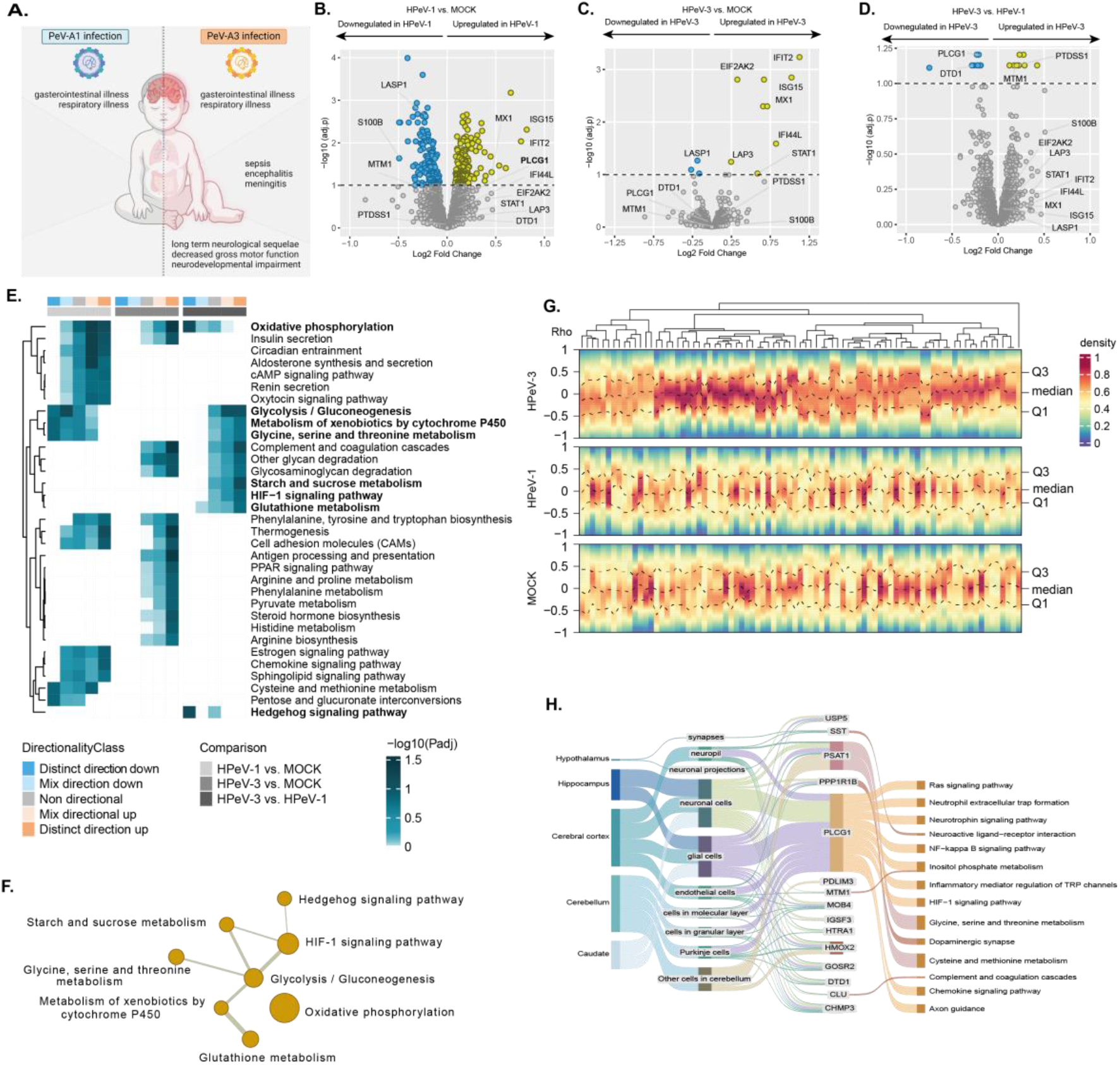
Proteome analysis of neural organoids shows proteins, pathways, and metabolic dysregulation unique for HPeV infection. **A**, differences in clinical presentation of HPeV-1 and HPeV-3 infection **B-D**, Volcano plots showing protein abundance in neural organoids in **B**, HPeV-1 vs mock **C**, HPeV-3 vs mock, and **D**, HPeV-3 vs HPeV-1 infection (adjusted *P* value < 0.1, Limma). The y-axis represents significance of protein abundance differences (negative log scale adjusted *P*-value) and the x-axis represents the fold change of the difference in abundance. **E-F**, Significantly regulated KEGG pathways linked to metabolism were analyzed where eight pathways are uniquely distinct for HPeV-3 infection in neural organoids. **E**, Pathway enrichment analysis results showing significantly regulated pathways (adjusted *P* value < 0.1, Limma) in the pair-wise proteomics analysis in neural organoids. This heatmap represents non, mixed, and distinct directionality (negative log scaled adjusted *P* value) of the pathways. The first column annotation denotes directionality of pathways and the second column denotes corresponding pair-wise comparison. **F**, Network visualization of within-pathway relationships between the eight significantly regulated pathways linked to metabolism unique for HPeV-3 infection. The node size represents the number of proteins annotated with this pathway and the line thickness is proportional the inter-pathway association. **G**, Density heatmap visualizing the correlation (Spearman’s rank correlation coefficient) between ISG/IFN related proteins and the metabolic DAPs in the proteome of HPeV-1, HPeV-3, or mock-infected organoids. Each column represents one ISG/IFN related protein, and dotted lines represent the mean values and quantiles. Data corresponds to two technical replicates (individual organoids) in three batches (independent experiments) of organoids. **H**, Sankey plot, depicting association between proteins uniquely abundant for HPeV-3 and respective tissue type, cell type and neural pathways. Data represents three technical replicates (individual organoids) for three batches (independent experiments) of organoids.

Next, we performed a pathway enrichment analysis to identify changes in pathways coupled with directionality of regulation. We found eight metabolic pathways to be uniquely distinct between HPeV-1 and HPeV-3 (figure 1E). Of these, two were distinctly downregulated (oxidative phosphorylation and hedgehog signaling pathway) and six were distinctly upregulated (glycolysis/ gluconeogenesis, glycine, serine and threonine metabolism, starch and sucrose metabolism, HIF-1 signaling pathway, and glutathione metabolism) in HPeV-3 infected neural organoids. Further analysis based on gene overlap showed interaction between the pathways suggesting a strong interplay of co-expressed proteins in HPeV-3 (figure 1F). As a distinct pattern for HPeV-3 infection in both innate immune and metabolic behavior was observed, we analyzed the relationship between these two host responses visualized in a heatmap (figure 1H). In contrast to the immunometabolic relationship in the mock neural organoids and HPeV-1 infected neural organoids, the heatmap representing HPeV-3 infected showed a disrupted relationship. This was evident from the high density around the Spearman’s rank correlation coefficient (ϱ) of zero (between Q1 and Q3), indicating a decrease or loss of correlation between the immune and metabolic pathways.

The cell-type specific protein expression maps identified both neuronal cells and glial cells as being associated with the DAPs unique to HPeV-3 infection(6). In addition, we mapped these DAPs to neurology-related pathways in the KEGG database(30), which showed links with multiple pathways, including complement and coagulation cascades and dopaminergic synapses (figure 1G). This analysis was also performed on the proteome dataset of HPeV clinical isolate infections, resulting in similar pathways to be uniquely abundant for infection with HPeV-3 (supplementary figure 1).

### HPeV-3 infection results in significantly higher extracellular glutamate and cytotoxicity in neural organoids

Among the DAPs unique to HPeV-3 infection were proteins involved in neurodevelopment and neuronal transmission, and have been associated with reactive astrocytes, excitotoxicity, and neurodegeneration (*e*.*g*., HTRA1, CLU, PLCy1, PSS1, DARPP32) in pathophysiological settings(31-37). In excitotoxic situations, the most abundant excitatory neurotransmitter, glutamate, is present at elevated levels in the synaptic cleft (figure 2A). Therefore, we performed glutamate assays on the supernatant of the neural organoids analyzed in our proteomics studies. Both HPeV-1 and HPeV-3 infection resulted in a significant increase in extracellular glutamate upon at 10dpi compared to the mock (figure 2B). A similar pattern was observed for lactate dehydrogenase (LDH), an important metabolic enzyme, at 10dpi upon HPeV-1 and HPeV-3 infection which is a direct indication of plasma membrane damage and cytotoxicity (figure 2D). Levels of Neurofilament Light (NfL), a biomarker for neuroaxonal damage, were also elevated in both HPeV-1 and HPeV-3 infected neural organoids at 10dpi compared to the mock (figure 2F), indicating neuronal damage following infection. Adjusting the levels of glutamate, LDH, and NfL to the viral load at 10dpi, the impact of HPeV-3 infection was significantly higher compared to HPeV-1 infection (figure 2C, E, G). To confirm this result, glutamate and LDH levels were measured using clinically relevant HPeV strains, revealing similar patterns (supplementary figure 2). This data indicates that HPeV-3 has a higher potential to trigger glutamate release or lead to glutamate accumulation and cytotoxicity in neural organoids. We, therefore, posit that HPeV-3 infection in neural organoids disrupts neuron-astrocyte communication, leading to increased extracellular glutamate and excitotoxicity. These results translate to clinical disease as HPeV-3 specifically can result in meningitis and encephalitis(18, 38).

**Figure 2.**
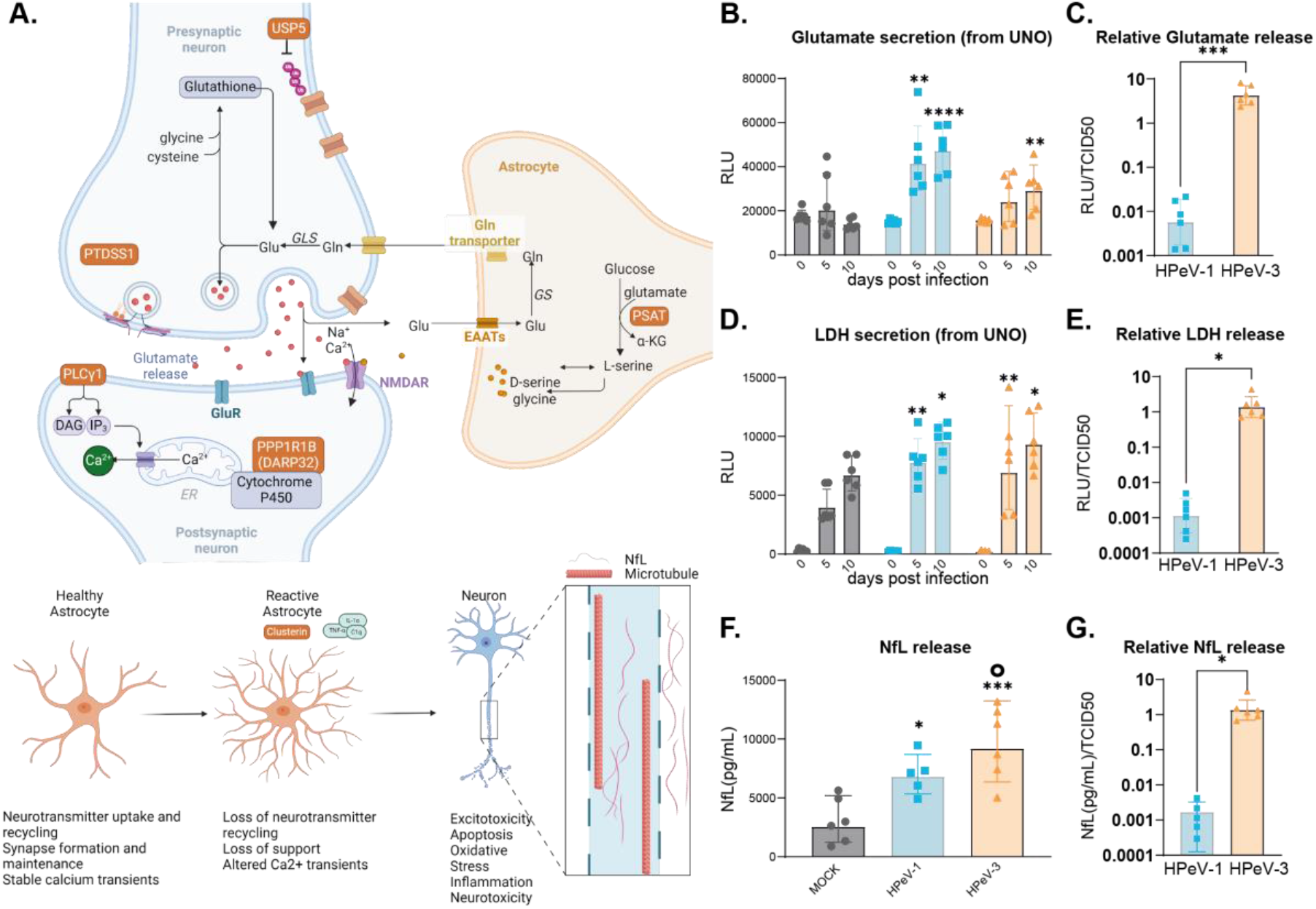
HPeV-3 infection results in significantly higher extracellular glutamate and cytotoxicity in neural organoids. **A**, Schematic representation of the proposed mechanisms of action of uniquely abundant proteins in HPeV-3 infection involving the glutamate cycle in a neuronal synapse. Glutamine is converted by glutaminase into glutamate, packed into vesicles, and released in a voltage dependent manner. After synaptic activation, the glutamate in the synaptic cleft is recycled predominantly by astrocytes and converted back to glutamine. Factors such as excess glutamate can trigger neurons to become reactive, which can induce loss of neuronal support and neurotoxicity. The proteins uniquely abundant in HPeV-3 infection are visualized in orange. **B-C**, Extracellular glutamate measured in the supernatant of HPeV-1, HPeV-3, or mock-infected neural organoids at 0, 5, and 10dpi, visualized as **B**, absolute data (RLU) or **C**, normalized to the amount of infectious viral particles (RLU/TCID50). **D-E**, Extracellular LDH measured in the supernatant of HPeV-1, HPeV-3, or mock-infected neural organoids at 0, 5 and 10dpi, visualized as **D**, absolute data (RLU) or **E**, normalized to the amount of infectious viral particles (RLU/TCID50). **F-G**, Extracellular NfL measured in the supernatant of HPeV-1, HPeV-3, or mock-infected neural organoids at 10dpi, visualized as **F**, absolute data (pg/mL) or **G**, relative to the viral load ((pg/mL)/TCID50). * significance compared to the mock of that respective dpi. ° represents significance of HPeV-3 vs HPeV-1. Data are represented as the geometric mean ± geometric standard deviation (SD) of three technical replicates (individual organoids) for three batches (independent experiments) of organoids. Statistical significance was analyzed (B,D,F) compared to mock using a one-way-ANOVA test with multiple comparisons or (C,E,G) comparing HPeV-1 to HPeV-3 using an unpaired t-test. *P value < 0.05; **P value < 0.01; ***P value < 0.001; ****P value < 0.0001. UNO; unguided neural organoid, LDH; Lactate dehydrogenase, NfL; Neurofilament Light, RLU; Relative Light Unit. TCID50; 50% tissue culture infectious dose.

### Astrocyte-neuronal co-cultures show significant increase in extracellular glutamate at early phases of HPeV-3 infection followed by an increase in CC3 expression

Upon HPeV-3 infection in neural organoids, there was a clear alteration of proteins and pathways involved in neural transmission. The proteomic data coupled with high extracellular glutamate and high levels of LDH and NfL after HPeV-3 infection pointed towards a disruption in the communication between the neurons and astrocytes. To understand the underlying mechanism and the initial cellular response upon infection, we inoculated 2D astrocyte-neuronal co-cultures with HPeV-1 and HPeV-3 at an MOI of 0.5 (figure 3A). Supernatant samples were collected to measure glutamate and LDH at 0-, 2-, 4-, 6-, 24-, 48-, and 72-hours post infection (hpi) and represented as a percentage of maximum release using a triton-X control (figure 3B,C). Mock-infection of the co-cultures did not result in a significant upregulation in glutamate at any of the time points (figure 3B). In HPeV-3 infected cocultures, increased glutamate secretion or lowered uptake was observed at early time points (0, 2, 4 and 6hpi) and was significantly higher compared to both mock and HPeV-1 infection. Although the levels of extracellular glutamate remained significantly higher up to 6 hours, a gradual decline could be observed over time. Furthermore, HPeV inoculation was not directly followed by a release of LDH after inoculation (figure 3C) suggesting that the released glutamate at these early time points was not due to membrane rupture or cell death. Subsequently, marker expression of β-tubulin, a structural neuronal protein, and cleaved caspase 3 (CC3), associated with initiation of apoptosis or necrosis, were measured in these cultures at 72hpi (figure 3 D,E,F). Quantification of β-tubulin and CC3 marker expression was performed and normalized to Hoechst as a readout of cell number. These results showed a significant upregulation of CC3 upon HPeV-3 infection, and a reduction in β-tubulin for HPeV-infected co-cultures compared to mock-infected co-cultures. Infection with clinical isolates of HPeV showed similar results (supplementary figure 3). Overall, these results corroborate the previous findings on HPeV infection of neural organoids and furthermore, suggest initiation of apoptotic cascades following glutamate excitotoxicity as a specific response to HPeV-3 infection.

**Figure 3.**
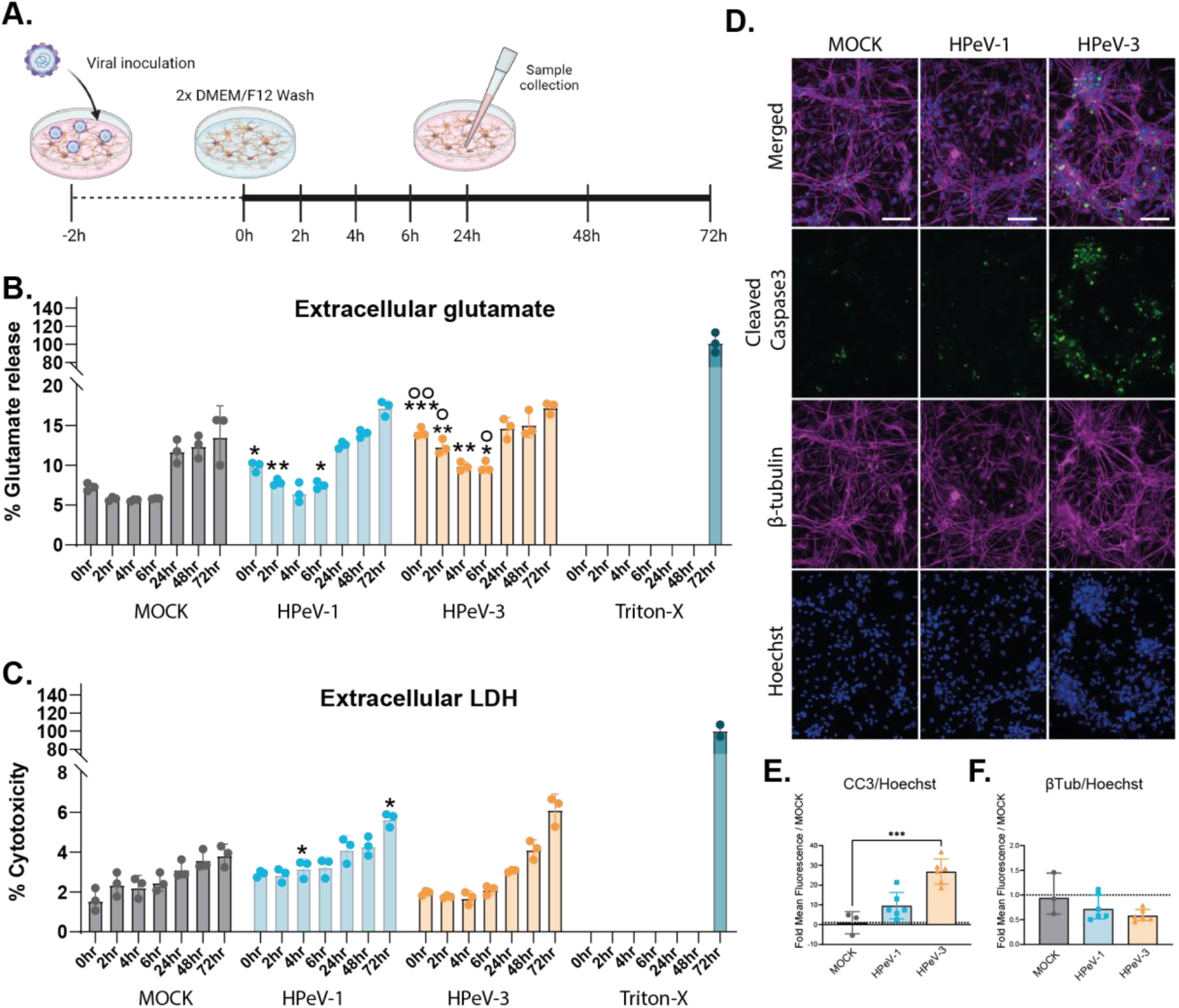
Astrocyte-neuronal co-cultures show significant increase in extracellular glutamate in the early phases of HPeV-3 infection and was followed by an increase in CC3 expression. **A**, Schematic representation of infection of the astrocyte-neuron co-culture with HPeV at a MOI of 0.5. **B-C**, Secretion levels at 0-, 2-, 4-, 6-, 24-, 48- and 72hpi measured in the supernatant of HPeV or mock-infected neural organoids as a percentage of maximum release control (Triton-X) of **B**, glutamate and **C**, LDH. Data corresponds to three technical replicates * represents significance of HPeV vs Mock, ° represents significance of HPeV-3 vs HPeV-1. **D-F**, Immunofluorescent images of HPeV-infected astrocyte-neuronal co-cultures stained for β-tubulin (magenta) and CC3 (green) at 72hpi. Scalebar represents 100µm. **E-F**, Quantification of mean fluorescent intensity of CC3 and β-tubulin relative to Hoechst as a fold change over MOCK. Data corresponds to the mean fluorescent intensity of representative images taken from multiple wells. Data are represented as the mean ± geometric standard deviation (SD) of three technical replicates. Statistical significance was analyzed compared to mock using a Kruskal–Wallis test with multiple comparisons,(E,F) Statistical significance was analyzed compared to mock using a one-way-ANOVA test with multiple comparisons *P value < 0.05; **P value < 0.01; ***P value < 0.001. LDH; Lactate dehydrogenase, CC3; Cleaved Caspase 3. hpi; hours post infection.

## Discussion

In this study, we identified potential metabolic factors involved in HPeV-3 CNS pathology using neural organoids. We assessed the presence of proteins and pathways unique for HPeV-3 infection, where, notably, many of the proteins and pathways were associated with glutamate metabolism. When dysregulated, these proteins can lead to hyperexcitability of neurons and, consequently, neurotoxicity. Moreover, we assessed whether HPeV-3 infection results in a dysregulation in the glutamate cycle by performing glutamate excretion and cytotoxicity assays. We validated our findings using an iPSC-derived neural-astrocyte co-culture. These insights advance our understanding of the metabolic factors contributing to HPeV-3 CNS pathology and underscore the potential importance of targeting glutamate metabolism in therapeutic strategies.

Infection in infants with HPeV-3 can cause neurological infections and long-term CNS pathology(15-17). Metabolic reprogramming can contribute to CNS pathology when initiated by viral infections(39, 40). The CNS exhibits a distinct metabolic profile where astrocytes and neurons interact extensively. Neurons are marked by their aerobic metabolism and high energy demand. In contrast, astrocytes, with their glycolytic metabolism, play a supporting role, including the regulation of neurotransmitters such as glutamate (41-43). Dysregulation of this neuronal-astrocyte crosstalk can lead to glutamate excitotoxicity, which is a key pathological factor in acute and chronic neurodegenerative diseases(22-24). Although little is known about excitotoxic neuronal injury in young children, defective glial glutamate re-uptake followed by apoptosis and programmed necrosis was reported, in which specifically activation of CC3 plays a major role(44, 45). Although conducted on cell lines, a previous study showed apoptosis, pro-apoptosis, and autophagy responses as primary causes for cell death due to HPeV-3 infection in neurons and astrocytes, including an increase in the level of CC3(46). While direct measures of glutamate levels in the CSF of patients with HPeV are not available, the clinical presentation involves symptoms consistent with excitotoxic damage, such as seizures and long-term neurological complications(15-17). Overall, metabolic reprogramming upon HPeV infection is interesting to investigate as a potential contributing factor for HPeV-3 neuropathology.

In the present study, we demonstrate 19 proteins to be uniquely altered following HPeV-3 infection, of which many were involved in CNS metabolism. Interestingly, among these were proteins essential for neurotransmission and maintenance of glutamatergic and dopaminergic neurons (e.g., HTRA1, CLU, PLCy1, PTDSS1, PSAT1, DARPP32). Dysregulation of many of these proteins has previously been associated with excitotoxicity of neurons due to increased cellular stress from elevated levels of reactive oxygen species (ROS) levels, and activation of the complement system(31-37, 47, 48). Moreover, the identified pathways uniquely upregulated for HPeV-3 infection are involved in neural plasticity, neurotransmission, and glutamate homeostasis. Furthermore, upon exhaustion of this glutamate cycle, glutathione can be compensatively increased to prevent decreased excitatory neurotransmission (49, 50) and was also found to be upregulated in our dataset. Lastly, upregulation of glycine, serine, and threonine metabolism, a pathway associated with dysregulation of glutathione, an increase of oxidative stress, loss of membrane potential and complement activation was uniquely upregulated for HPeV-3 in our results (51, 52). These findings further confirm the involvement of glutamatergic neurons, astrocytes, excitatory neurotransmitters, ROS, and the complement system in the pathological process of HPeV-3.

Another key finding was a dysregulated link between interferon-related proteins and metabolism that was unique to HPeV-3 infection in neural organoids. Previous studies report an interdependency between interferon signaling pathway and cellular metabolism(1, 4, 5, 53). We previously reported that HPeV-3 replication was reduced by a host cell response involving JAK/STAT signaling pathways and ISG transcription(6). In contrast to HPeV-3, HPeV-1 appears to circumvent activation of the JAK/STAT pathway, avoiding a subsequent innate inflammatory response, and therefore, maybe also preventing metabolic dysregulation. This observed immunometabolic dysregulation unique for HPeV-3 is of interest as a potential contributing factor in the development of neurologic disease.

Based on our results in neural organoids and published data on the involvement of DAPs unique for HPeV-3 in excitotoxicity, we hypothesized a potential dysfunction in the neurotransmission cycle following HPeV-3 infection. In physiological conditions, extracellular substrates as a result of synaptic neuronal activity (i.e., glutamate), are repleted by activated astrocytes to restore extracellular homeostasis (43, 54). The abundant extracellular glutamate unique to HPeV-3 infection in neural organoids, therefore, indicates re-uptake of glutamate by astrocytes may not have been sufficient. The corresponding high levels of toxicity markers such as extracellular LDH and NfL confirm that this extracellular glutamate concentration reaches toxic levels. Plasma and CSF levels of NfL, a cytoskeletal protein specifically expressed in neurons, are used as a state-of-the-art biomarker for neuroaxonal damage and is associated with encephalomyelitis and cognitive impairment. As elevated NfL levels were previously shown to correlate to long-term impaired cognitive performance following viral neuroinflammation(55, 56), the high levels in the supernatant of neural organoids infected with HPeV-3 could correspond to this clinical presentation.

To understand the underlying mechanism and the initial cellular response upon infection at early time points, we also investigated glutamate release upon HPeV infection in astrocyte-neuronal co-cultures. The astrocyte-neuronal co-cultures infected with HPeV-3 showed significant increase in extracellular glutamate at early phases of infection when LDH release was not yet present. This indicates HPeV-3 triggers neurotransmitter release *via* a different route other than membrane disruption or cytotoxicity. Interestingly, the observed increase in glutamate release within one hour of viral infection suggests that it is worth investigating whether the entry of the virus itself, besides viral replication, triggers glutamate release and inhibits subsequent uptake. Further analysis of marker expression showed that at 72dpi there was a significant increase in CC3 specific to HPeV-3 infection in the astrocyte-neuronal co-cultures. Overall, this data points towards HPeV-3 triggered glutamate excitotoxicity as a potential mechanism contributing to initiation of apoptotic cascades leading to neuronal toxicity.

In summary, based upon our results we hypothesize that CNS infection with HPeV-3 can result in neuronal toxicity and neuronal cell death via glutamate excitotoxicity and loss of astrocyte-neuronal communication. This is consistent with other CNS pathologies caused by viruses (e.g. West Nile Virus infections(57), human immunodeficiency Virus-1(58) and SARS-CoV-2 (59)), including long-term pathologies. These insights are pivotal in identifying metabolic targets for therapeutic interventions in severe neonatal neurological infections caused by HPeV-3.

## Methods

### Cell lines and virus strains

Virus stocks were generated on cell lines, as described previously(6). In short, human colorectal adenocarcinoma (HT-29) cells (HTB-38™ ATCC), rhesus monkey kidney cells (LLCMK2, provided by the Municipal Health Services, the Netherlands), African green monkey kidney cells (Vero, provided by the National Institute of Public Health and the Environment, RIVM, the Netherlands) were used for virus culture. Lab adapted virus strains used were HPeV-1 Harris and HPeV-3 152037. Clinical isolates (<10 passages in cell lines) used were HPeV-1 52967, HPeV-1 51067, HPeV-3 178608, and HPeV-3 51903. The TCID50 of the viral stocks was calculated using the Reed and Muench method(60). Genome Sequences of these strains are available on GenBank (accession numbers BankIt: OR886056-OR886061).

### Human induced pluripotent stem cell culture

Human induced pluripotent stem cell (hiPSCs) were obtained from WiCell (IMR90-4/WISCi004, WiCell) and cultured in mTeSR^+^ medium (STEMCELL Technologies) supplemented with 1% (v/v) Pen-Strep on human laminin 521 (Biolamina)-coated culture ware. Subculturing was performed weekly using ReLeSR (STEMCELL Technologies) according to the manufacturers protocol with the addition of 10 µM Y-27632 Rho Kinase (ROCK) inhibitor (Cayman Chemical Company) on the day of passaging. Daily medium changes were performed and cultures were visually assessed for differentiation before passaging. The maintenance and subsequent experiments using hiPSCs were performed in accordance with relevant guidelines and regulations. Cells were routinely tested for mycoplasma.

### Generation and infection of unguided neural organoids

Generation and infection of unguided neural organoids (UNOs) was performed as previously reported(6). In short, UNOs were generated using human induced pluripotent stem cells (hiPSCs) (IMR90-4/WISCi004-B, WiCell) and STEMdiff™ Cerebral Organoid kit from STEMCELL™ Technologies. At day 67 neural organoids were inoculated with 10^5^ TCID50 per mL of HPeV lab-adapted strains and clinical isolates. After 2 hours of inoculation at 37 °C and 5% CO_2_, UNOs were washed three times with phosphate buffer saline (PBS, Lonza), and moved to a freshly coated 48-well plate with 500 µL of Maturation Medium (STEMCELL™ Technologies). Collection and replenishment of medium was performed at day 1, 3, 5, 7, and 10dpi. Organoids were harvested at 10dpi for proteomic analysis. This experiment was performed three times independently, using individually generated neural organoid batches, and within each experiment two technical replicates per condition.

### Generation of an astrocyte-neuronal co-culture

hiPSC derived neural progenitor cells (NPCs) were generated using the embryoid body protocol and STEMdiff SMADi Neural Induction Kit (STEMCELL Technologies). From these NPCs, both astrocytes and neurons were generated usig the STEMdiff Astrocyte Differentiation and Maturation kit, and the STEMdiff Forebrain Neuron Differentiation and Maturation kit, respectively. The astrocytes were maintained in long term culture for 52 days and analysed for marker expression (GFAP and S100B). hiPSC-derived neurons were cultured for 24 days and analysed for β-tubulin expression (. Astrocytes and neurons were co-cultured to model relevant cell-cell interactions *in vitro* in a 2D culture. Tissue culture treated 48-well plates (Sarstedt 83.3923) and Ibidi µ-Slide 18 Well (Ibidi 81816) were coated at 37°C for 2 hours with 15 µg/mL Poly-L-ornithine (PLO, P4957, Sigma) in phosphate buffer saline (PBS, Lonza) followed by two washes with PBS and 5 µg/mL Laminin (L2020, Sigma) at 37°C for 2 hours. Neurons were dissociated according to the STEMCELL protocol and plated at a density of 5 × 10^4^ cells/cm^2^ on the coated cultureware. The following day, astrocytes were dissociated according to STEMCELL protocol, and plated at a density of 1 × 10^5^ cells/cm^2^ on top of the neuronal culture (2:1 ratio) in STEMdiff Astrocyte Maturation Medium. After 24 hours, the medium was removed and replenished with STEMdiff Forebrain Neuron Maturation Medium. The co-culture was maintained at 37°C and 5% CO_2_, with full medium changes using Forebrain Neuron Maturation Medium performed every 2-3 days for 7 days before used for subsequent experiments. Before infection, mono-cultures and co-culture were analyzed for quality control purposes by immunofluorescent staining (supplementary figure 4).

### Infection of astrocyte-neuronal co-cultures

On the day of infection, the co-cultures were inoculated with MOI 0.5 of the different virus stocks or mock-infected and incubated at 37°C for 2 hours. Samples were taken for back titrations of viral inoculums and comparable inoculation titers were confirmed using TCID50. The wells were carefully washed twice using DMEM/F12 and replenished with 400 µL STEMdiff Forebrain Neuron Maturation Medium per well. Collection of medium was performed at 24, 48, and 72 hpi where samples from the same wells were taken and stored at -70°C until further processing.

### Immunofluorescence staining

Astrocytes and neuron monoculture or co-cultures on Ibidi µ-Slide 18 Well (Ibidi 81816) were washed with PBS and fixed at 72hpi using 4% (v/v) formaldehyde (Sigma Aldrich) in PBS for 20 minutes at room temperature (RT) followed by three more washes with PBS. Before immunofluorescence staining, the wells were submerged in blocking solution (10% (v/v) Sea block Blocking Buffer (Thermo Fisher Scientific), 1% (v/v) Triton X-100 (Sigma) in PBS) for 2 hours at RT. All antibodies were diluted in 1:1 blocking buffer: PBS solution. Primary antibodies were added and incubated overnight at 4°C. After three washes with PBS, the secondary antibodies and 1:1000 Hoechst (Thermo Fisher Scientific) staining were added and incubated at RT for 1h. Samples were washed three times with PBS and quenched using ReadyProbes Tissue Autofluorescence Quenching kit (Invitrogen, kit) for five minutes followed by another PBS wash. Antibody dilutions and details are provided in Supplementary table 1. Slides were imaged using a Leica TCS SP8-X microscope and Leica LAS AF Software (Leica Microsystems), EVOS M5000 microscope (Thermo Fisher Scientific) or MICA Wide Fied/confocal Fluorescence Microscope (Leica Microsystems) and Leica LAS-X Software (Leica Microsystems).

### Image analysis

Representative images from a minimum of four replicate wells were taken for image analysis. Quantification of marker expression was performed by quantifying the mean fluorescent intensity of each marker using ImageJ 1.50I, and normalized to Hoechst as a readout of cell number. Marker expression was shown as a fold change over MOCK-infected controls.

### Bioinformatics analysis

Detailed bioinformatics methodologies used to process the proteomics data was provided in the previous publication(6). Briefly, R package Linear Models for Microarray Data (Limma) v3.50.0(61) was used to find differentially abundant proteins and pathway enrichment analysis was performed using R package Piano v2.18.0 (nPerm = 1000, geneSetStat = mean, and signifMethod = geneSampling)(62). KEGG pathway gene-sets obtained from enrichr libraries(62) was used for the pathway enrichment. Pathways belonging to the categories metabolism, environmental information processing, organismal systems were used. Cell type specific protein expression information was obtained from the human protein atlas (version 23.0 and Ensembl version 109)(63). The data consists of expression profiles for proteins in human tissues based on immunohistochemisty using tissue micro arrays. Heatmap was generated using R package ComplexHeatmap v2.18.0(64). Volcano plots and sankey plots were generated using R packages ggplot2 v3.5.1 and ggsankey v0.0.9 respectively

### Glutamate secretion assay

Glutamate-Glo™ Assay (Promega) was used, according to the manufacturer’s instructions, for the detection of glutamate in supernatant samples of both neural organoid samples, and 2D astrocyte-neuronal co-culture. In short, 5 μL of supernatant was stored in 95 μL of PBS, and stored at -70°C until further processing. The assay was performed by transferring 25 μL of the sample in PBS to a white opaque-bottom 96-well plate. Glutamate detection reagent was added to the sample at a 1:1 ratio (25 μL) and incubated at RT in the dark for 1h. Luminescent signal was measured using an H1 Synergy plate reader (BioTek). Data was normalized to relevant controls.

### LDH secretion assay

LDH-Glo™ Cytotoxicity Assay (Promega) was used according to manufacturer’s protocol in neural organoid supernatant and supernatant of the 2D astrocyte-neuronal co-culture. In short, 5 μL of supernatant was stored in 95 μL of LDH Storage buffer (200 mM Tris-HCl pH7.3, 10% v/v glycerol, 1% w/v BSA) and stored at -70°C until further processing. The assay was performed by adding 50 μL LDH detection reagent to 50 μL of the sample in LDH storage buffer in a black clear bottom 96-well plate incubated at RT in the dark for 1h. Luminescent signal was measured using an H1 Synergy plate reader (biotech). Data was normalized to relevant controls. For the 2D co-culture experiment, maximum LDH Release Control was included, where triplicate wells were treated with 2 μL of 10% Triton® X-100 per 100 μL to mock-infected 2D astrocyte-neuron co-culture for 10–15 minutes before collecting the samples for LDH detection. This control was used to calculate the percentages of cytotoxicity and glutamate release.

### Neurofilament Light (NfL) assay

Supernatant of neural organoids infected with HPeV-1, HPeV-3, or mock-infected (10dpi) was diluted 1:100. NfL concentrations were measured using an in-house developed Simoa assay using a Single Molecule Array (Simoa) NF-lightTM ® Advantage Kit run on automated HD-X Analyzer (Quanterix, Lexington, MA, USA) (65).

### Data visualization and statistical analysis

All statistical analysis other than LC-MS/MS-based quantitative proteomic analysis was performed using GraphPad Prism 8 (GraphPad Software Inc.). Data represents two technical replicates from three independent experiments using three independent organoid batches unless otherwise stated. All statistical analytical tests performed for each analysis are indicated in the corresponding figure legend.

## Data availability

Sequencing results are available on GenBank (accession numbers BankIt: OR886056-OR886061). The mass spectrometry proteomics data have been deposited to the ProteomeXchange Consortium via the PRIDE partner repository with the dataset identifier PXD047238.

## Acknowledgements

All schematic representations were created with BioRender.com. We wish to acknowledge Charlotte E. Teunissen and Johannes A. Heijst for their contribution and technical assistance on the Neurofilament Light Assays. We also thank the Municipal Health Services and the National Institute of Public Health, and the Environment (RIVM) for the supply of the cell lines. Finally, we acknowledge AII Work visit grant for facilitating a work visit between the Amsterdam UMC and Karolinska Institutet. This research was funded by the European Union’s Horizon 2020 Research and Innovation Programme under the Marie Sklowdowska-Curie Grant Agreement OrganoVIR (grant 812673, KW/DP),GUTVIBRATIONS (grant 953201, KW/DP), and the PPP Allowance (Focus-on-Virus, DP/KW) made available by Health Holland, Top Sector Life Sciences & Health, to the Amsterdam UMC, location Amsterdam Medical Center to stimulate public–private partnerships. UN acknowledges support from the Swedish Research Council grants 2021-00993 (UN) and 2021-01756 (UN) and Karolinska Institute Consolidator Grant (2-117/2023) (UN).

## Author Information

### Contributions

Conceptualization: P.C., A.S., U.N,. Data curation: P.C., I.G.-R., and A.T.A., Formal analysis: P.C., I.G.-R., A.T.A. Funding acquisition: D.P., K.W, A.S., and U.N. Investigation: P.C., I.G.-R., A.T.A., Methodology: P.C., I.G.-R., A.T.A., and R.V.-S. Project administration: K.W., D.P., A.S., and U.N. Supervision: R.V.-S., U.N., D.P., A.S., and K.W. Validation: P.C., I.G.-R. Visualization: P.C., A.T.A., U.N., and A.S. Writing of the original draft: P.C., A.T.A., U.N., and A.S. Writing, review, and editing: P.C., I.G.-R., A.T.A., R.V.-S, K.W., D.P., A.S., and U.N. As described by CRediT-Contributor Roles Taxonomy (casrai.org). All authors contributed to the article and approved the submitted version.

## Supplemental Information

Document S1 - Supplemental figures 1-4 and Supplementary table 1

