## Supplementary Information for "Parechovirus-3 infection disrupts immunometabolism and leads to glutamate excitotoxicity in neural organoids"

Supplementary figure 1

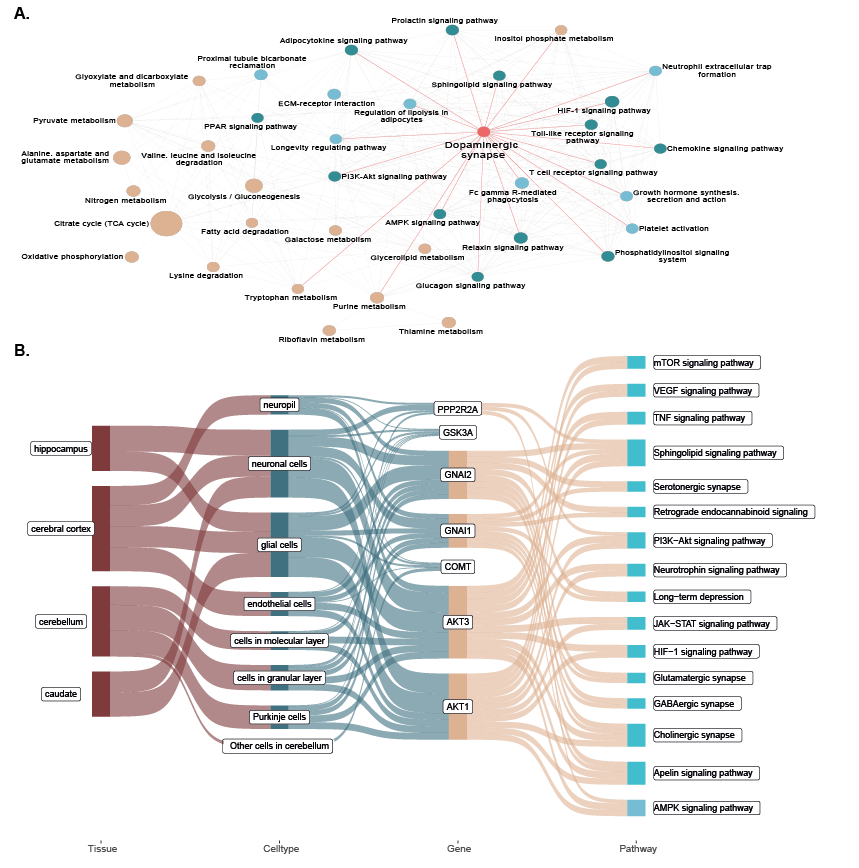

**Supplementary Figure 1. Dopaminergic synapse pathway enriched upon HPeV3 clinical isolate infections in neural organoids at 10dpi.** **A,** Network visualization of significantly enriched pathways (padj < 0.2) in HPeV-3 infection compared to HPeV-1. Nodes represent pathways and edge represent gene overlap between a pair of pathways. Node size is relative to negative log10 scaled adjusted pvalues obtained from enrichment analysis. Association between neural pathway and other signaling and metabolic pathways based on gene overlap are highlighted.

**B**, Sankey plot showing brain tissue and cell type specific expression information of significantly regulated (pvalue < 0.05) proteins in HPeV-3 compared to HPeV-1 that are part of dopaminergic synapse pathway and the association of them with other signaling pathways. Data are represented are three technical replicates (individual organoids) for three batches (independent experiments) of organoids.

Supplementary Figure 2

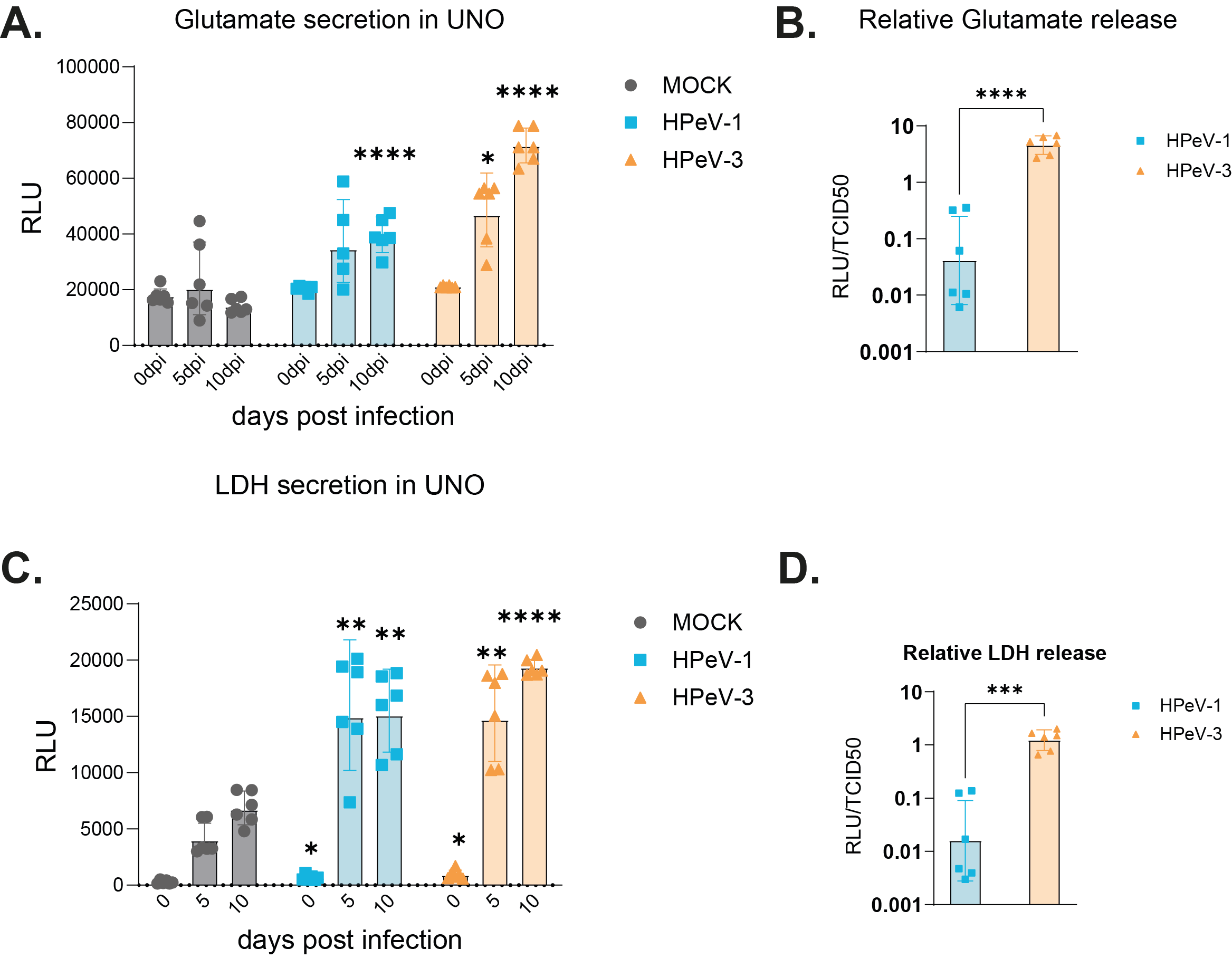

**Supplementary Figure 2. HPeV-3 infection using clinical strains results in significantly higher glutamate release and cytotoxicity in neural organoids. A-B,** Glutamate secretion measured in the supernatant of HPeV-1 clinical strains, HPeV-3 clinical strains, or mock infected neural organoids at 0, 5 and 10dpi, visualized as A**,** absolute data (RLU) or **B,** normalized to the amount of infectious viral particles (RLU/TCID50). **C-D,** LDH secretion measured in the supernatant of HPeV-1, HPeV-3 or mock infected neural organoids at 0, 5 and 10dpi, visualized as **C,** absolute data (RLU) or **D,** normalized to the amount of infectious viral particles (RLU/TCID50). Data are represented as the geometric mean±geometric standard deviation (SD) of three technical replicates (individual organoids) for three batches (independent experiments) of organoids. Statistical significance was analyzed (B,D,F) compared to mock using a one-way-ANOVA test with multiple comparisons or (C,E,G) comparing HPeV-1 to HPeV-3 using an unpaired t-test. *P value<0.05; **P value<0.01; ***P value<0.001; ****P value<0.0001. UNO; unguided neural organoid, LDH; Lactate dehydrogenase, RLU; Relative Light Unit. TCID50; 50% tissue culture infectious dose

Supplementary Figure 3

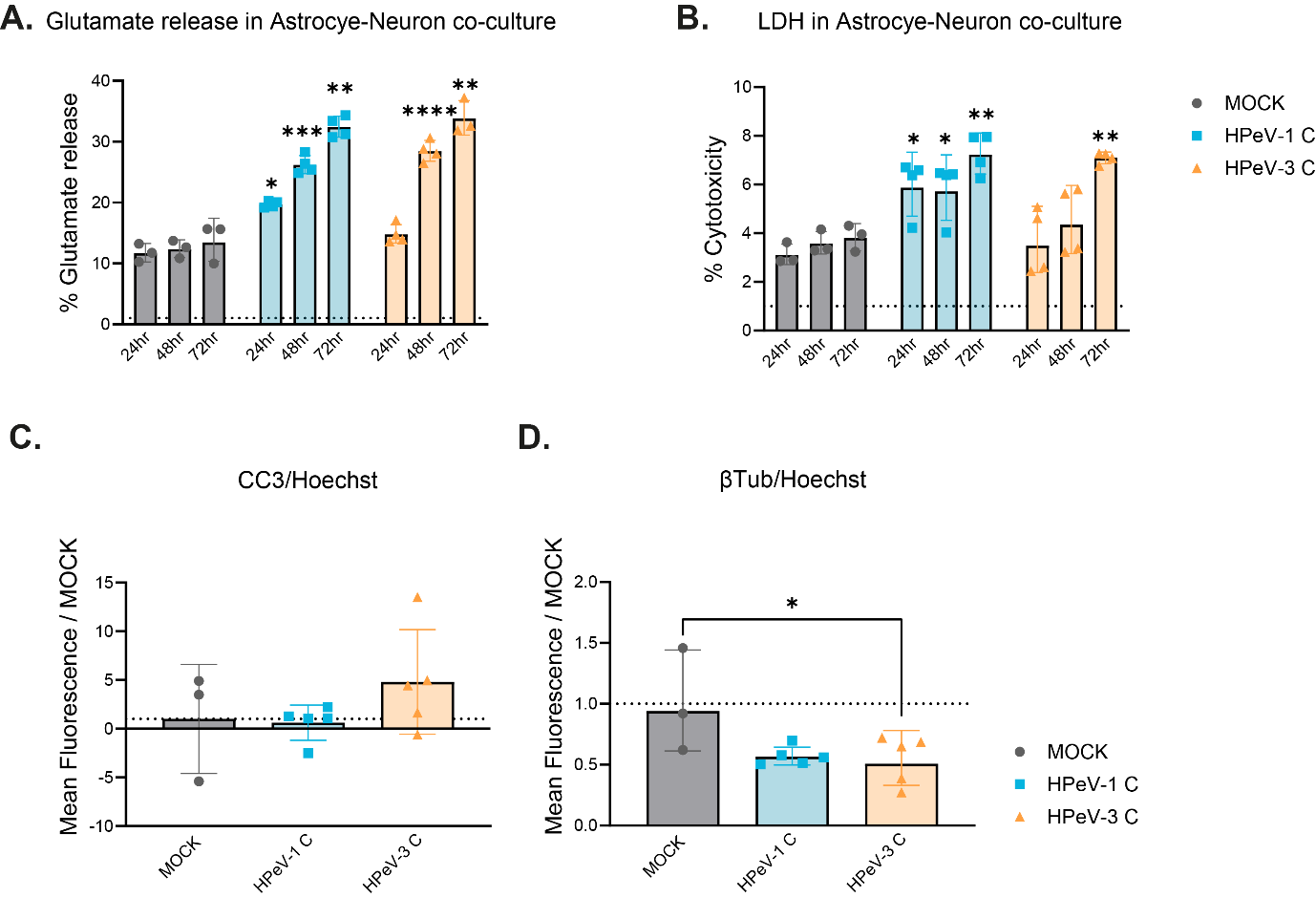

**Supplementary Figure 3. Extracellular glutamate and LDH levels and marker expression in Astrocyte-neuronal co-cultures upon HPeV infection using clinical isolates**

**A-B**, Secretion levels at 24-, 48- and 72hpi measured in the supernatant of HPeV or mock infected neural organoids as a percentage of maximum release control (Triton-X) of **b,** glutamate and **c,** LDH. Data corresponds to three to four technical replicates * represents significance of HPeV vs Mock, ◦ represents significance of HPeV-3 vs HPeV-1. **C-D,** Quantification of immunofluorescent images of HPeV infected astrocyte-neuronal co-cultures stained for β-tubulin (magenta), and CC3 (green) at 72hpi relative to Hoechst as a fold change over MOCK. Data corresponds to the mean fluorescent intensity of representative images taken from multiple wells. Data are represented as the mean±geometric standard deviation (SD) of three technical replicates. **A-D,** Statistical significance was analyzed compared to mock using a Kruskal–Wallis test with multiple comparisons,(E,F) Statistical significance was analyzed compared to mock using a one-way-ANOVA test with multiple comparisons *P value<0.05; **P value<0.01; ***P value<0.001; ****P value<0.0001. LDH; Lactate dehydrogenase, CC3; Cleaved Caspase 3. hpi; hours post infection.

Supplementary Figure 4

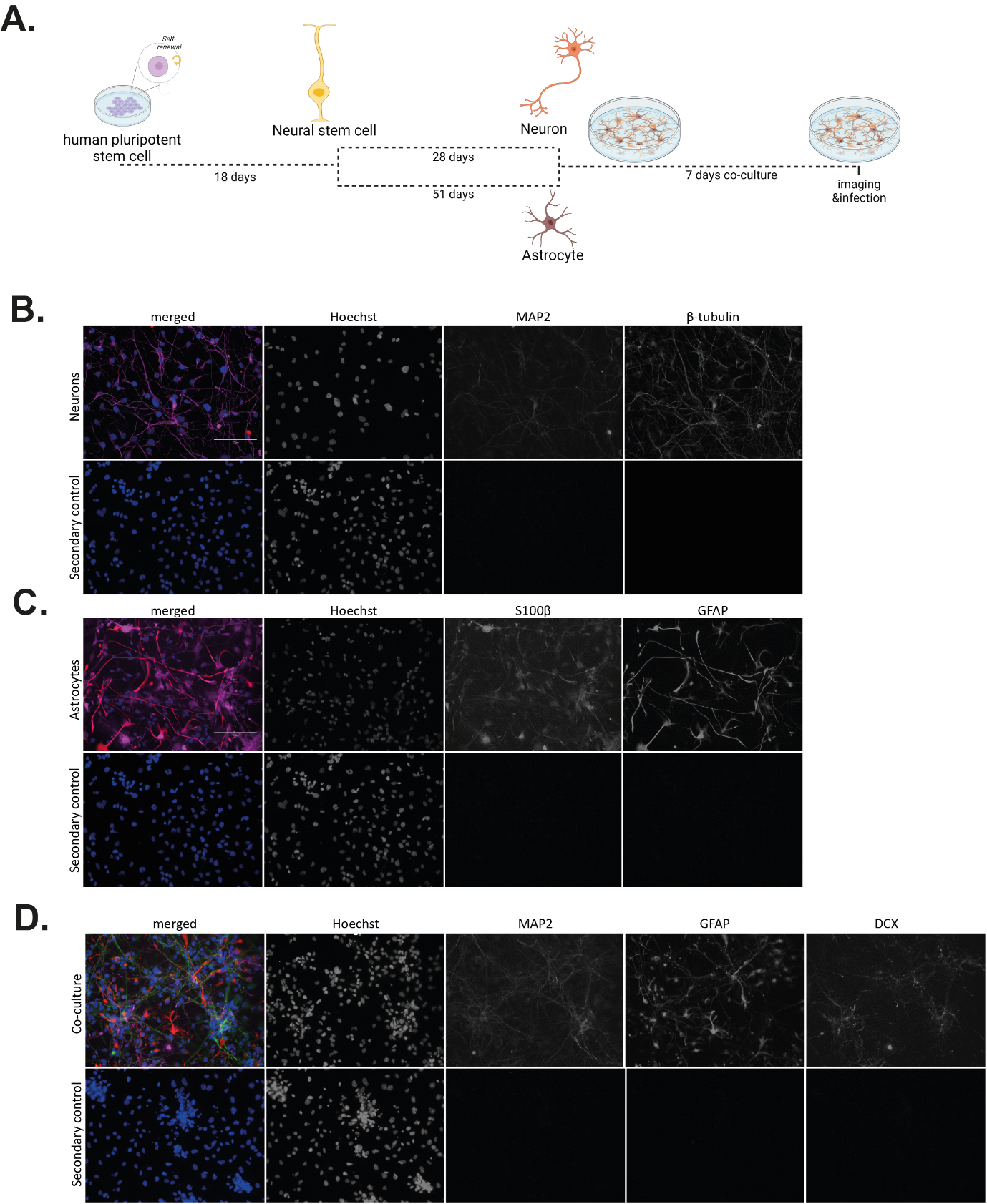

**Supplementary figure 4. Validation of stem cell derived astrocytes and neurons used to generate an astrocyte-neuronal co-culture.** a, Schematic representation and timeline of the generation of astrocytes and neurons and co-culture. b, validation of neuronal marker expression by using MAP2, and β-tubulin. c, Validation of Astrocyte marker expression by using GFAP and S100β. d, Marker expression of co-cultured astrocytes and neurons after 7 days in culture (MAP2, GFAP, DCX).

**Supplementary table 1. Primary and secondary antibodies used**

| **Antigen (abbreviation)** | **Origin** | **Dilution factor** | **Company** | **Catalog number** |
| --- | --- | --- | --- | --- |
| Glial fibrillary acidic protein (GFAP) | Goat | 1:500 | Abcam | Ab53554 |
| Microtubule-associated protein 2 (MAP2) | Mouse | 1:500 | Thermo Fisher | MA5-12826 |
| β-Tubulin (β-Tub) | Chicken | 1:200 | Abcam | ab41489 |
| S100β | Rabbit | 1:500 | Abcam | ab52642 |
| Doublecortin (DCX) | Rabbit | 1:500 | Thermo Fisher | 48-1200 |
| dsRNA clone rJ2 | Mouse | 1:60 | Sigma-Aldrich | MABE1134 |
| Cleaved Caspase 3 (CC3) | Rabbit | 1:200 | Cell Signaling Technologies | 96615 |
| **Secondary Antibody (origin)** | **Host** | **Dilution factor** | **Company** | **Catalog number** |
| Anti mouse Alexa Fluor 546 | Donkey | 1:750 | Thermo Fisher | A10036 |
| Anti rabbit Alexa Fluor 488 | Donkey | 1:750 | Thermo Fisher | A21206 |
| Anti chicken Alexa Fluor 647 | Donkey | 1:750 | Thermo Fisher | A78952 |
| Anti mouse Alexa Fluor 488 | Donkey | 1:750 | Thermo Fisher | A21202 |
| Anti rabbit Alexa Fluor 647 | Donkey | 1:750 | Thermo Fisher | A31573 |
| Anti chicken Alexa Fluor 546 | Donkey | 1:750 | Thermo Fisher | A11056 |
